# Deciphering the role of histo-blood group antigens in bovine rotavirus C infection

**DOI:** 10.1101/2025.03.10.642353

**Authors:** Noemi Navarro-Lleó, Jonathan Ramírez-Cárdenas, Francisco Paredes-Martínez, Sonia Serna, Roberto Gozalbo-Rovira, Juan C. Muñoz-García, Niels C. Reichardt, Patricia Casino, Jesús Angulo, Jesús Rodríguez-Díaz, Javier Buesa

**Affiliations:** Departament de Microbiologia, Facultat de Medicina i Odontologia, Universitat de València, 46010 Valencia, Spain; Instituto de Investigaciones Químicas (CSIC—Universidad de Sevilla), 41092 Sevilla, Spain; Departamento de Bioquímica y Biología Molecular, Universitat de València, Burjassot, 46100 Valencia, Spain; Instituto Universitario en Biotecnología y Biomedicina (BIOTECMED), Universitat de València, Burjassot 46100 Valencia, Spain; Glycotechnology Laboratory, Center for Cooperative Research in Biomaterials (CIC biomaGUNE), Basque Research and Technology Alliance (BRTA), 20014 Donostia-San Sebastián, Spain; INCLIVA. Instituto de Investigación Sanitaria del Hospital Clínico de Valencia, 46010 Valencia, Spain; CIBER-BBN, 20014 Donostia-San Sebastián, Spain; CIBER de Enfermedades Raras (CIBERER-ISCIII), Spain

## Abstract

Rotaviruses (RVs) are the main cause of viral diarrhea among infants, small children, and the young of many animal species. Histo-blood group antigens (HBGAs) are potential RV receptors and glycan composition on mucous surfaces influences host susceptibility and cross-species virus transmission. RVs exhibit genotype-dependent glycan binding and differences are due to sequence modifications in the VP8* domain of the spike protein VP4. Nevertheless, the molecular bases for this genotype-dependent glycan specificity, especially in non-A RVs, are not thoroughly understood. This study delves into how genotypic variations configure a novel binding site in the VP8* of a bovine P[3] rotavirus species C (RVC) strain to recognize H type-2 antigen (H2) and its precursor N-acetyl-lactosamine (LacNAc) using glycan binding assays, crystallography, and STD NMR. Results reveal a specific interaction of bovine P[3] RVC VP8* with H2 and LacNAc, more strongly with the latter. In the P[3] RVC VP8*-H2 interaction, the N-acetyl glucosamine moiety displays significant interaction, while galactose participates moderately and fucose binds weakly. Moreover, the bovine VP8* structure, resolved at 3 Å, shows specific structural features which differ from human RVC, as it contains two additional β-strands (β1 and β2) contributing to β-sheet2 and conformational changes that widens the cleft to allow different carbohydrate binding modes. These subtle changes in both sequence and structure explain the H2 precursor recognition, which is ubiquitous in human neonate intestine and in human and bovine milk, providing insights into P[3] RVC tropism and its potential zoonotic transmission.

**Author Summary:** Rotavirus C (RVC) represents an emerging pathogen with the ability to infect both humans and animals. It is widely acknowledged that host glycobiology plays a crucial role in determining susceptibility to RVs. Consequently, a better understanding of the interactions between RVs and carbohydrates is essential to know how the virus causes infection and to develop effective preventive strategies. Recent findings have unveiled the capability of the human RVC genotype P[2] to recognize the type A antigen through the VP8* spike protein although little is known about animal RVC strains. In this study, we describe the capacity of bovine P[3] RVC strain to bind H2 and LacNAc, both HBGAs. We also offer the structural explanation for the glycan differential recognition between human and bovine RVC strains. Our findings offer valuable insights into RVC attachment to host cells and its potential implications for species barriers.

## Introduction

Rotaviruses (RVs) belong to the *Rotavirus* genus within the *Sedoreoviridae* family, and are the predominant enteric viral agents causing acute gastroenteritis in both young humans and animals worldwide [1], resulting in over 240,000 deaths in children under 5 annually [2].

The RV virion is composed of three concentrically arranged protein layers surrounding the viral genome: the core (VP2), the inner capsid (VP6), and the outer capsid (VP7, VP4) [3]. Based on the VP6 sequence, RVs are classified into 9 species/groups designated alphabetically from A to D and F to J [4–6], with *Rotavirus alphagastroenteritidis* (RVA) being the predominant cause of diarrheal disease in humans [2]. The prevalence of infections caused by non-A RV species remains unknown. Notably, species RVA, RVB, RVC, and RVH, can infect both humans and animals remarking their zoonotic potential. In contrast, RVD, RVF, RVG, RVI, and RVJ only cause animal infections [7].

RVs have a segmented genome composed of 11 double-stranded RNA segments encoding six structural (VP1-4, VP6, and VP7) and five or six non-structural proteins (NSP1-NSP5/6) [3]. RVs can be classified by a binary classification system [1] based on the VP4 and VP7 proteins of the outer capsid, which determines G genotypes (dependent on VP7, a glycoprotein) and P genotypes (dependent on the protease-sensitive VP4 protein). The rotavirus surface spike protein, VP4, is cleaved by trypsin into two non-covalently bonded polypeptides, a head subunit (VP8*) and a stem subunit (VP5*). While both subunits stay connected on the virion surface, the galectin-like domain VP8* is responsible for the recognition of host cell glycans as receptors facilitating initial cell attachment and infection [8].

RVA has demonstrated the ability to recognize cell surface glycans as essential ligands or receptors, playing a critical role in rotavirus attachment and cell entry [9–11]. Numerous animal RVAs recognize the internal sialic acids (SAs) of sialoglycoconjugates, including gangliosides [12], while some human RVAs specifically recognize terminal SAs [13]. Additionally, many other human RVAs exhibit binding affinity for non-sialylated glycoconjugates like histo-blood group antigens (HBGAs), HBGA precursors, and/or mucins [9,14,15]. HBGAs are carbohydrates expressed on the surfaces of red blood cells, gastrointestinal epithelial cells, and mucosal secretions [10]. Their synthesis is governed by a set of glycosyltransferases known as enzymes A, B, FUT2, and FUT3. Genetic polymorphisms in these enzymes lead to variability in antigen expression among individuals influencing the interaction, susceptibility, or resistance to RVs that bind HBGAs [16].

*Rotavirus tritogastroenteritidis* or species C rotavirus (RVC) has emerged as a pathogen causing gastroenteritis in neonatal piglets and is increasingly reported in both pigs and humans [17–19]. RVCs have also caused infections in cows, ferrets, and dogs [20–23]. Global reports indicate RVC infections in humans across North and South America, Asia, Africa, Europe, and Oceania [22]. Beyond human RVC, numerous seroepidemiological studies in pigs have shown the circulation of these viruses in swine herds across different countries for many decades, with seroprevalence rates ranging from 58% to 100% [24]. Furthermore, at least two studies from the USA demonstrated remarkably high RVC prevalence in suckling piglets [18,25].

In contrast to RVA, receptors or attachment factors associated with the adhesion and infection of RVC have received comparatively less attention. Limited research has explored the role of SAs and HBGAs in rotavirus infections caused by these non-A species. Nevertheless, a recent study [26] demonstrated that the VP8* of human RVC G4P[2] recognizes type A HBGAs through a mechanism distinct from that used by RVA. Similarly, another study revealed that a trisaccharide glycan, Galα1-3Galβ1-4Glc, containing a terminal α-Gal, was recognized by multiple RVA/RVC genotypes, suggesting a shared evolutionary origin linked to the synthesis of HBGAs in animals [27]. Therefore, further exploration of the diverse interactions between RVs and host glycans holds the potential to enhance our understanding of RV epidemiology, prevalence, possible zoonotic transmission, and the development of novel prevention strategies.

To investigate whether the bovine genotype of RVC exhibits the ability to bind to HBGAs similarly to the human genotype, the VP8* protein from the bovine P[3] RVC genotype was produced recombinantly in *Escherichia coli* with a N-terminal GST tag. Its interaction with a panel of synthetic HBGAs was subsequently examined, revealing a robust binding affinity with H2 (Fuc-α1,2-Gal-β1,4-GlcNAc) and its non-fucosylated precursor, LacNAc (Gal-β1,4-GlcNAc). Additionally, it was established that the N-acetyl-glucosamine moiety of H2 directly interacted with the VP8* protein. Furthermore, the crystal structure of bovine RVC VP8* was determined, revealing two twisted β-sheets separated by a cleft resembling the galectin-fold which showed structural differences to the human RVC VP8* strain. The bovine RVC VP8* showed two additional small strands, β1 and β2, contributing to a six-stranded β-sheet2 similar to the S-face of galectin. Also, it showed a flexible long loop βJ-βK instead of a β-hairpin, and βK was connected to β2 via a short loop, lacking this region three residues that are present in human RVC, thus, widening the cleft between β-sheets. Finally, docking studies were performed, uncovering a novel glycan-binding pocket specific to bovine P[3] RVC genotype.

## Results

### Bovine P[3] RVC VP8* recognize terminal Galβ1-4GlcNAc and GalNAcβ1-4 GlcNAc epitopes

Our investigation started by elucidating the glycan binding specificity of bovine P[3] RVC VP8*, based on the crucial role of the VP8* domain interacting with host receptors [10,15,28]. The recombinant VP8* protein was purified as a GST fusion protein and was employed to incubate a glycan microarray containing a library of 155 different structures prepared chemoenzymatically as previously described [29,30]. The library contains a collection of N-glycan structures as found on glycoproteins of mammalian, invertebrate and plant origin along with smaller O-glycan structures and structural elements of larger glycan structures [31]. The bovine VP8* exhibited significant binding signals to sixteen glycans from the library, including both N-glycan and O-glycan structures all displaying terminal LacNAc (Galβ1-4GlcNAc) or LacdiNAc (GalNAcβ1-4GlcNAc) residues (Fig 1). The GST-bovine VP8* protein exhibited binding signals with typical cores of O-glycosylation, specifically O-glycosylated mucins (glycan numbers GL138, GL139, GL142, and GL143) (Fig 1).

**Fig 1.**
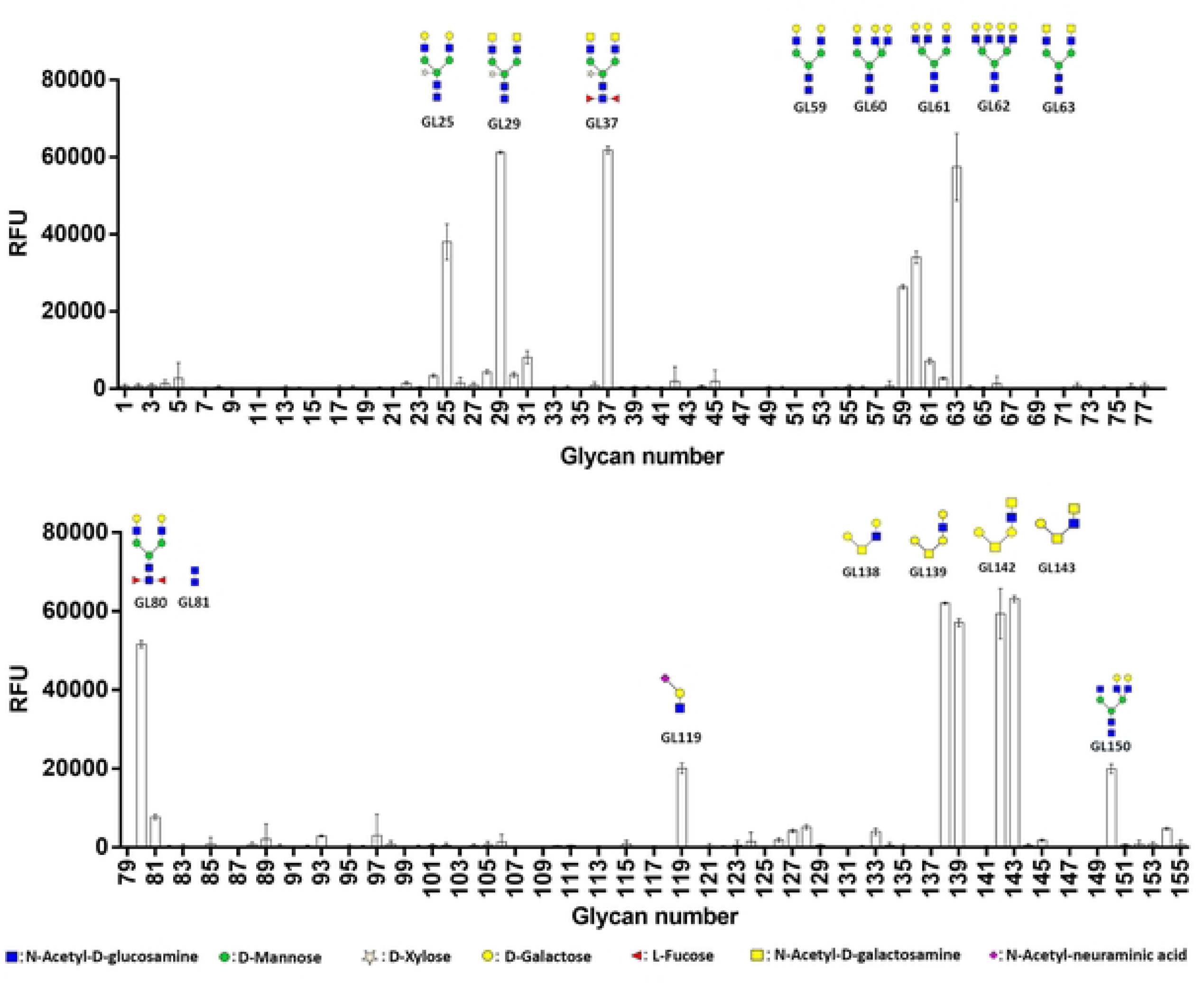
Bovine P[3] RVC GST-VP8* protein incubation on the glycan microarray. Histograms representing mean RFU (Relative Fluorescence Units) values after incubation with bovine P[3] RVC GST-VP8* protein. Each histogram represents the average RFU values of four replicate spots with the standard deviation of the mean. The structure of the glycans showing signal binding is represented according to the standard Symbol Nomenclature for Glycans (SNFG) v.1.5. (See S1 Fig, for the glycan structures included in the microarray screening).

### The VP8* domain of the bovine P[3] RVC genotype interacts with both the H2 antigen and its precursor LacNAc

To further validate the observed ligand binding selectivity, we carried out glycan interaction studies using synthetic oligosaccharides representing HBGAs and GM3 (S1 Table). Bovine VP8* protein demonstrated notable interaction signals with two HBGAs, consisting of H2 antigen and its precursor LacNAc (Fig 2), both containing Galβ1-4GlcNAc epitope. Furthermore, the results showed that the VP8* protein of bovine RVC recognized specifically the H2 antigen and LacNAc, although binding to the second antigen was 7 times stronger (Fig 2) (P value =0.0006). However, minimal or no signals were detected for the remaining assayed glycans. In summary, our data clearly indicate that the VP8* domain of bovine P[3] RVC specifically recognize LacNAc and H2 HBGAs.

**Fig 2.**
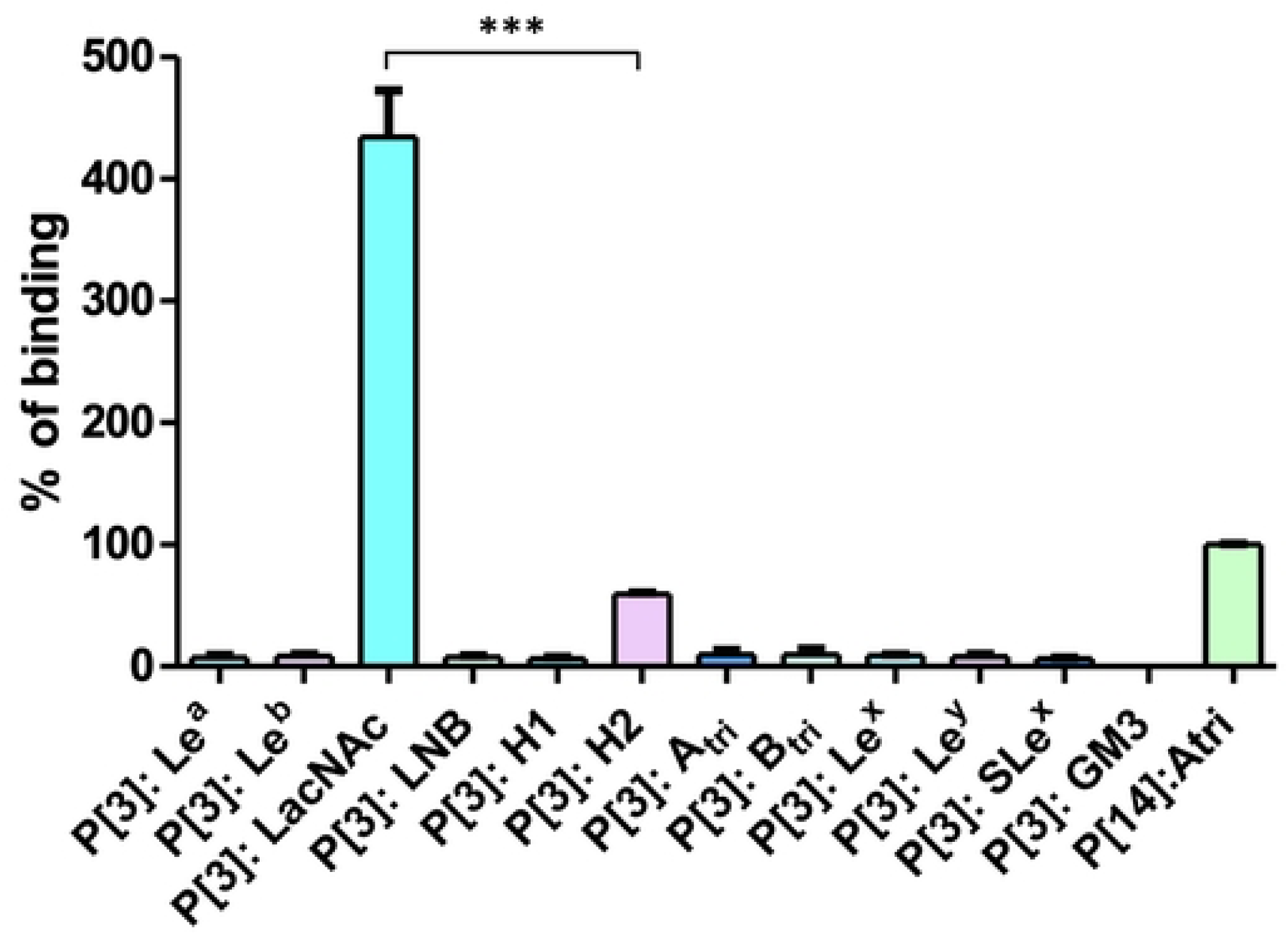
ELISA-like glycan binding assay of bovine P[3] RVC VP8* genotype to a panel of biotinylated sugars. RV-glycan binding (*y* axis) was measured in units of fluorescence (F.U.) normalized to positive control the RVA P[14] GST-VP8* recombinant protein. GST was included as a negative control, and its fluorescence signal was subtracted for each interaction involving bovine VP8*. P value < 0.001 (***). Bars indicate the standard error of the mean.

### Binding epitope mapping of H-type 2 to bovine P[3] RVC VP8*

To obtain structural details and identify the moiety of H2 antigen interacting with the bovine RVC VP8* protein we obtained their binding epitope mappings by Saturation Transfer Difference (STD) NMR spectroscopy experiments. The NMR method is based on the saturation transfer from the protein to the bound ligand, which is transferred further to the free ligand where the signal is detected due to rapid exchange at the binding site. Ligand protons close to the protein binding site receive substantial saturation and exhibit greater effects in STD NMR compared to protons more exposed to the solvent [32,33], so that the relative level of saturation received by the ligand protons can be translated into a ligand-binding epitope map, depicting the key contacts of the ligand for the interaction with the protein in the binding site.

All performed STD NMR experiments showed clear signals confirming the binding of H2 antigen with bovine VP8*. In this study, two binding epitope mappings were obtained: one corresponding to the antigen with the reducing GlcNAc in α configuration (Fig 3A), and that with the GlcNAc in β configuration (Fig 3B), as they produced two sets of ^1^H NMR signals for the reducing GlcNAc ring and only one set of signals for the remaining galactose and fucose rings. After analyzing the integral values for the signals corresponding to the anomeric protons of α and β-GlcNAc, a higher concentration of the α conformation in solution was inferred. The different concentrations of both anomers can lead to biased estimations of the binding epitope mappings so, to avoid overestimations in the binding epitope of the lower populated β-anomer, a precise comparison between the existing concentrations of α and β anomers was carried out using the law of mass action (ligand concentration 2 mM, protein concentration 50 µM, and an estimated dissociation constant of 500 µM, consistent with this type of protein-carbohydrate interaction). In this way, to obtain appropriate binding epitopes, we considered the differences in the concentrations of both anomers by weighting the measured STD intensities by the proportion of β anomer in comparison to the α one, so that the STD_0_ values for the β-GlcNAc protons were weighted according to equation 1.

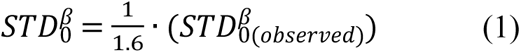

**Fig 3.**
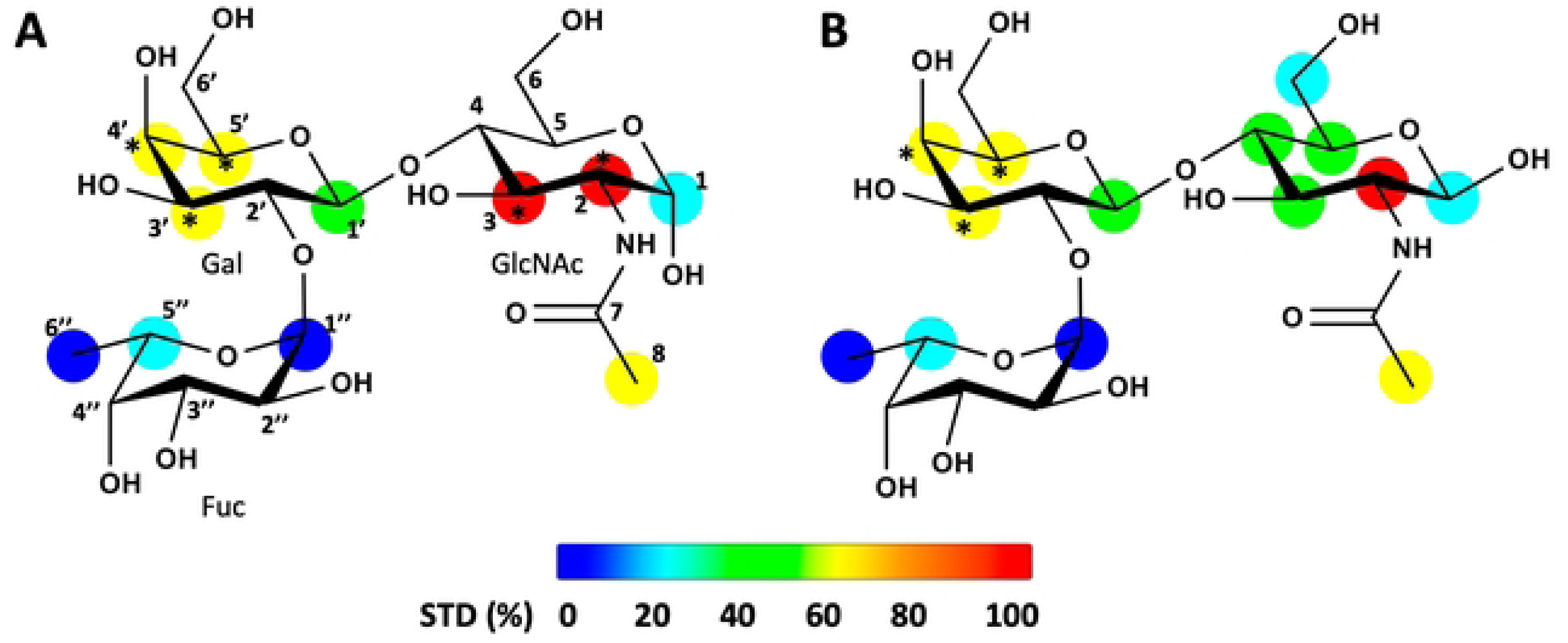
STD NMR binding epitope mappings of H2 antigen to bovine P[3] RVC VP8*. **A** Binding epitope maps of the H2 antigen Fucα1-2Galβ1-4GlcNAcα and **B** Fucα1-2Galβ1-4GlcNAcβ. Protein saturation was achieved by irradiation at -2 ppm. Normalized STD-NMR intensities are shown in different colored spheres as represented in the colored legend bar. STD responses are only indicated for protons that could accurately be measured. Asterisks indicate averaged STD values due to overlapping. STD NMR: saturation transfer difference NMR, Fuc: fucose, Gal: galactose, GlcNAc: N-acetyl-glucosamine.

The results indicate a slight preference for interaction with the α-configuration of the β-GlcNAc ring, although the observed binding epitope mapping is rather similar in both cases (Fig 3). Globally, there is a significant preference for the reducing GlcNAc ring, a reduced contribution of the galactose ring, and a much lower contribution of the fucose ring in the recognition of the trisaccharide by the VP8* protein.

### Crystal structure of bovine P[3] RVC VP8*

To understand the molecular basis of the interaction between bovine RVC VP8* and H2, we carried out crystallization studies with VP8* alone and in the presence of H2. Crystals were obtained just for bovine RVC VP8* (residues comprised from E64 to R228) which diffracted X-rays at 3 Å resolution in the space group P2_1_2_1_2_1_ (Table 1). The asymmetric unit of the crystal contained two molecules (Fig 4A), each one formed by two twisted antiparallel β-sheets; β-sheet1 formed by six strands (βA, βL, βB, βI, βJ and βK) and β-sheet2 formed by six strands (βC, βD, βG, βH and two small strands β1 and β2) (Fig 4B). The presence of two additional β-strands in β-sheet2 can reproduce the S-face of galectins [34], however, there is a β-hairpin, formed by βE and βF, inserted between βD and βG which sits in the groove of the β-sheet2 blocking the sugar binding site at S-face observed in galectins (Fig 4B and S2A Fig). These two additional β-strands (β1 and β2) and the β-hairpin insertion are also present in RVA VP8* (PDB: 6H9W) (S2A Fig) [15]. Structural comparison between the two molecules of RVC VP8* found in the asymmetric unit showed high similarity (0.4 Å RMSD for 153 residues) but, conformational changes were observed in the loop β1-βB and loop βJ-βK located in β-sheet1 (Fig 4B), thus, indicating flexibility for this β-sheet, which resembles the F-face of galectins (S2B Fig). This fact could indicate a tendency of this face to discriminate between glycans, as the loop βJ-βK seems to adopt different conformations in VP8* contributing to the cleft configuration [35]. Meanwhile, β1 and part of the loop β1-βB can be involved in interactions with VP5* according to the structure of VP4 spike [36].

**Fig 4.**
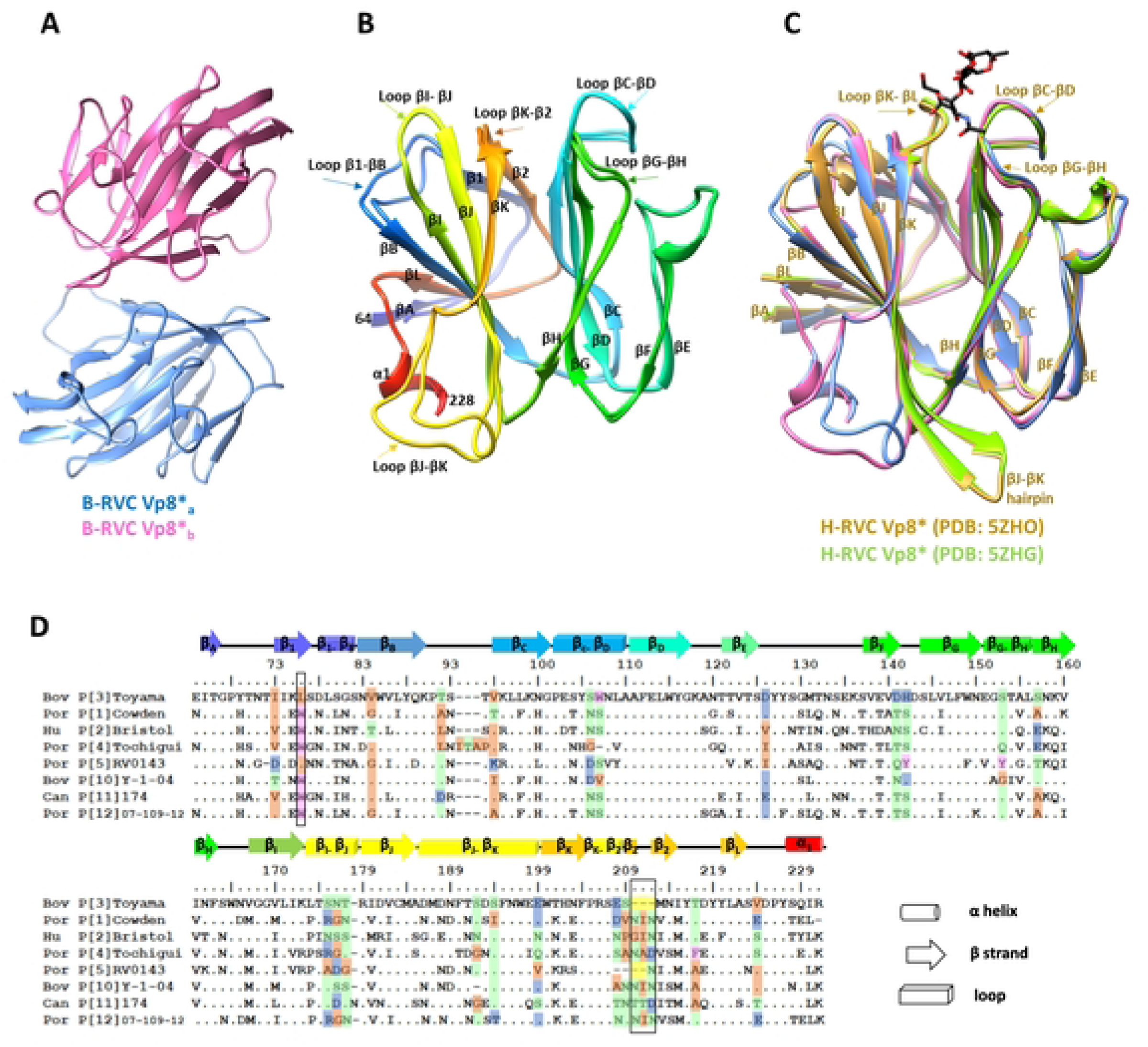
Structure of VP8* of bovine P[3] RVC (B-RVC). **A** The asymmetric unit of the crystal contained two molecules: a (in blue) and b (in pink). **B** Secondary structure comparison of the two molecules found in the asymmetric unit, in rainbow coloring, formed by two twisted β-sheets; β-sheet1 formed by six strands (βA, βL, βB, βI, βJ and βK) and β-sheet2 formed by six strands (βC, βD, βG, βH and two small strands β1 and β2) shows differences in the conformation of β1-βB loop connecting β1 and βB strands as well as between βJ-βK loop connecting βJ and βK strands. **C** Superposition of the two B-RVC VP8* molecules with VP8* from human RVC (H-RVC) in the absence (PDB: 5ZHG in green colour) and presence (PDB: 5ZHO in yellow color) of antigen A trisaccharide (color stick representation). Nomenclature of β-strands has been adopted from H-RVC VP8* published in [26]. **D** Sequence alignment based on the structure of the galectin-like domain of VP8* proteins of different RVC P genotypes. The alignment was constructed using Clustal Omega and the residues less conserved were colored, assigned non-polar aliphatic in orange, polar uncharged in green, charged in blue, aromatic in pink, and sulfur-containing in purple. The less conserved positions predicted by PRALINE are highlighted by boxes and the secondary structure of B-RVC P[3] is shown.

**Table 1.** Data collection and refinement statistics for the obtained structures.

|  | P[3] RVC VP8* |
| --- | --- |
| **Data collection** |  |
| Space group | P2<sub>1</sub> 2<sub>1</sub> 2<sub>1</sub> |
| Cell dimensions |  |
| *a*, *b*, *c* (Å) | 61.49 64.37 76.94 |
| $\alpha$, $\beta$, $\gamma$ (°) | 90.00 90.00 90.00 |
| Resolution (Å) | 74.94-3.00 (3.18-3.00) |
| No. reflections | 83626 (13907) |
| *R*<sub>sym</sub> or *R*<sub>merge</sub> | 0.354 (1.049) |
| *R*<sub>pim</sub> | 0.143 (0.420) |
| *I*/σ*I* | 7.7 (3.1) |
| Completeness (%) | 99.90 (99.70) |
| Redundancy | 12.9 (13.5) |
| **Refinement** |  |
| *R*<sub>work</sub>/*R*<sub>free</sub> | 0.24/0.29 |
| No. atoms |  |
| Protein | 2572 |
| Water | 51 |
| B-factors (Å<sup>2</sup>) |  |
| Protein | 35.69 |
| Water | 20.26 |
| R.m.s deviations |  |
| Bond lengths (Å) | 0.002 |
| Bond Angles (°) | 0.578 |
| PDB Code | 9EN4 |

A search for structural homologs with the DALI server indicated high structural similarity with human RVC VP8* [26], both in the absence (PDB: 5ZHG) and presence of A trisaccharide antigen (PDB: 5ZHO) (0.7 Å RMSD for 134 residues), however, we observed several differences in their secondary structure compared to bovine RVC (Fig 4C). First, the bovine RVC contained two additional β-strands (β1 and β2) forming part of β-sheet2 (Fig 5A) where the end of β1 corresponds to a helix turn in human RVC, and thus, major changes in the conformation of the loop that connects with βB are observed (Fig 5A). Second, the flexible long loop βJ-βK in bovine RVC is a part of the long beta hairpin in human RVC (Fig 4C and Fig 5A). Third, in bovine RVC, βK is connected to β2 via a short loop, lacking this region three residues that are present in human RVC. Interestingly, this region is part of the binding pocket for the A trisaccharide antigen in human RVC, thus, the shortening of this region in bovine RVC has widened the cleft between both β-sheets possibly changing the specificity for glycan binding [26] (Fig 4C and Fig 5B). Surprisingly, a sequence variation between bovine and human RVC seems to link the configuration of β1 and loop β1-βB with the shortening of loop βK-β2 and β2. In this way, in bovine RVC there is a Leu at the end of β1 beginning of loop β1-βB, which is a Trp in human RVC as part of a helix turn (Fig 5A). The side chain of Trp lies on the short loop βK-β2 of bovine RVC, thus, only in the absence of a Trp in this position can the shortening of loop βK-β2 exist. Lastly, the C-terminal end of bovine RVC was longer than human RVC containing one helix turn (Fig 4C and Fig 5A). Overall, despite the structural homology with human RVC, VP8* molecules from bovine RVC show specific structural features that contribute to the widening of the cleft, such as the short loop βK-β2 linked to β1 and loop β1-βB as well as to the flexible long loop βJ-βK (Fig 5B), which could allow different carbohydrate binding modes.

**Fig 5.**
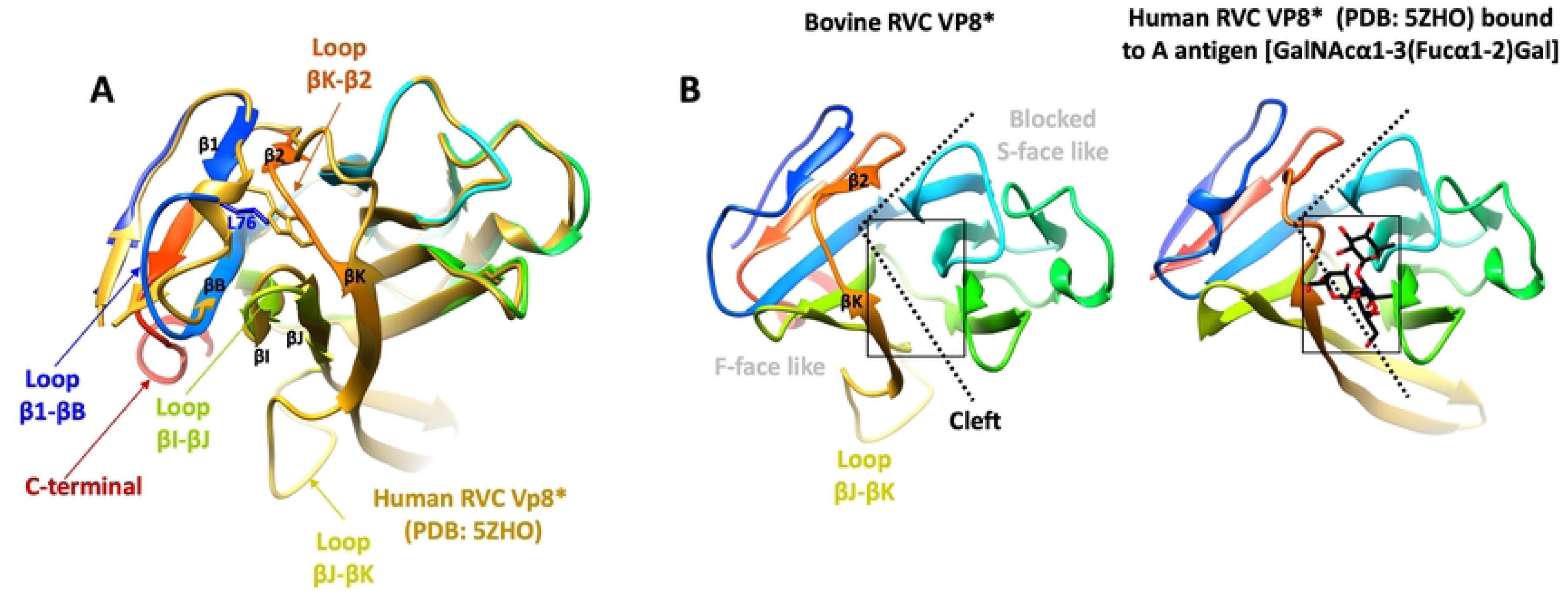
Comparison of the structures of bovine RVC VP8* and human RVC VP8* (PDB: 5ZHO). A) Superposition of both structures, bovine RVC VP8* (in rainbow color) and human RVC VP8* (in gold) shows several structural differences in β-sheet2: the presence of β1 and β2 in bovine RVC VP8*, presence of Leu in loop β1-βB instead of Trp and conformational changes in this loop, loop βI-βJ, loop βJ-βK and loop βK-β2. B) Superposition of both structures, shown separated for clarity, and in rainbow color. In human RVC VP8*, the A antigen is shown as sticks and framed by a square in both structures to indicate the position. Also, the cleft between β-sheet2 and β-sheet1 is denoted by two dashed lines, which demonstrate that it is wider in bovine RVC VP8*.

### Structure and sequence analysis of RVC VP8*

We conducted sequence alignments to find correlations for our structural changes in different human and animal genotypes of the RVC VP8* galectin-like domain proteins (residues 64 to 231). The overall sequence identity between them was 66.5% but non-synonymous amino acid substitutions were identified (Fig 4D). We observed that at the end of β1/beginning of loop β1-βB all genotypes contained Trp, as in the structure of human RVC, except in porcine P[5] and in our bovine P[3] RVC VP8* structure which contained Leu (Leu76; Fig 5A). As we mentioned previously, the absence of Trp widens the cleft for glycan binding, an effect that can be observed upon modelling the structures of the different genotypes with AlphaFold (S3 Fig). Also, the loop βC-βD which is involved in glycan binding at the structure of human VP8* RVC, contains Trp (W107) in bovine P[3] VP8* but this position was Ser in human P[2], porcine P[1], P[5], P[12] and canine P[11] while Val for bovine P[10] (Fig 4D). Additionally, the loop βI-βJ showed sequence variability between genotypes. Interestingly, the less conserved position was found between the short loop βK-β2 (Arg206-Glu208) and β2 (Ser209-Asn211) as it contained a deletion of three residues in the bovine RVC P[3] that was not present in other genotypes, except for porcine P[5] (Fig 4D). As mentioned previously, this deletion, identified in the structural comparison between human P[2] and bovine P[3] VP8*, is concomitant with the presence Leu, instead of Trp, at the end of β1/beginning of loop β1-βB. Hence, just the bovine P[3] and porcine P[5] genotypes show either the deletion or presence of Leu which contributes to the widening of the cleft between β-sheets. This fact may result in modifications of the glycan recognition region at bovine P[3], and possibly in porcine P[5] as well, that may account for the switch in specificity from type A antigen to LacNAc and H2 antigens.

### Molecular docking and molecular dynamics simulations reveal a novel glycan binding pocket for P[3] bovine RVC

To obtain a 3D model of the molecular complex between bovine RVC VP8* and H2 antigen, molecular docking calculations were performed for the α and β anomers of the H2 antigen using different VP8* conformations (generated from MD (molecular dynamics) simulations of the *apo* protein). Docking of the H2 antigen in both α and β configurations produced about 1000 docking poses for each anomer. These were subsequently assessed against experimental STD NMR data using a software recently developed *in-house*, called RedMat, that enables the fast calculation of theoretical STD NMR binding epitopes mappings from both static (docking simulations) and dynamic (MD simulations) 3D models [37]. This allowed us to perform model validation using the NOE *R*-factor as the metric to assess the goodness between theoretical and experimental binding epitopes and, thus, to filter the 3D models of the VP8*-H2 complex in agreement with the experimental data (i.e., those NOE *R*-factors below 0.3). A total of seven validated models were obtained.

To analyse the stability in solution of the validated 3D models, we performed 500 ns of MD simulations. For the H2 antigen in β configuration, only one of the validated models exhibited ligand stability at the recognition site throughout the entire MD trajectory (S4A Fig), while the simulations of the other models showed the ligand dissociating from the protein within the first 100 nanoseconds. RedMat was employed to monitor the evolution of the NOE *R*-factor along the stable simulation, yielding values below 0.3 in the majority of frames and, hence, validating this dynamic 3D model against the experimental STD NMR data (S4B Fig). Overall, this highlights the importance of combining RedMat calculations with the assessment of model stability along an MD simulation for the robust validation of 3D models of protein-ligand complexes.

The same procedure was followed for the H2 antigen in α configuration, and a 3D model was obtained where the orientation of the ligand was very similar to the validated model of the β configuration. Subsequent 500 ns of MD simulation confirmed ligand stability at the recognition site, as shown by the orientation of the ligand (RMSD; S5A Fig), and the NOE R-factor during the trajectory (S5B Fig). Higher NOE R-factor values were observed for the α compared to those obtained for the β configuration, although no significant changes in ligand orientation were observed (S5A Fig and S1 Video). This increased NOE R-factor for the α anomer simulation can be attributed to proton H2, which despite receiving the highest saturation according to the experimental binding epitope (100%), is averaged in the analysis with H3 due to peak overlapping in the ^1^H NMR spectrum (Fig 3). This leads to a miscalculation of the STD_0_ of H2 in the α anomer, therefore introducing some bias into the whole binding epitope since the STD_0_ of all other protons are normalized against H2. However, this does not occur for the β configuration of the H2 ligand.

Since the α configuration of H2 is the major anomer, the frame of the α H2 MD simulation showing the smallest NOE R-factor has been selected as the 3D model best representing the molecular recognition between VP8* and the H2 antigen (Fig 6A). MD simulation shows that VP8* recognizes H2 antigen, interacting with GlcNAc via Ala110 and Glu208 and Gal and Fuc moiety via Arg206 and Leu109, respectively (Fig 6A). It is important to highlight that the H2 antigen is placed in the VP8* cleft interacting with the βK-β2 loop via both residues Arg206 and Glu208 (Fig 6A). In addition, the theoretical binding epitope of this model showed great agreement with the experimental one (Fig 6B).

**Fig 6.**
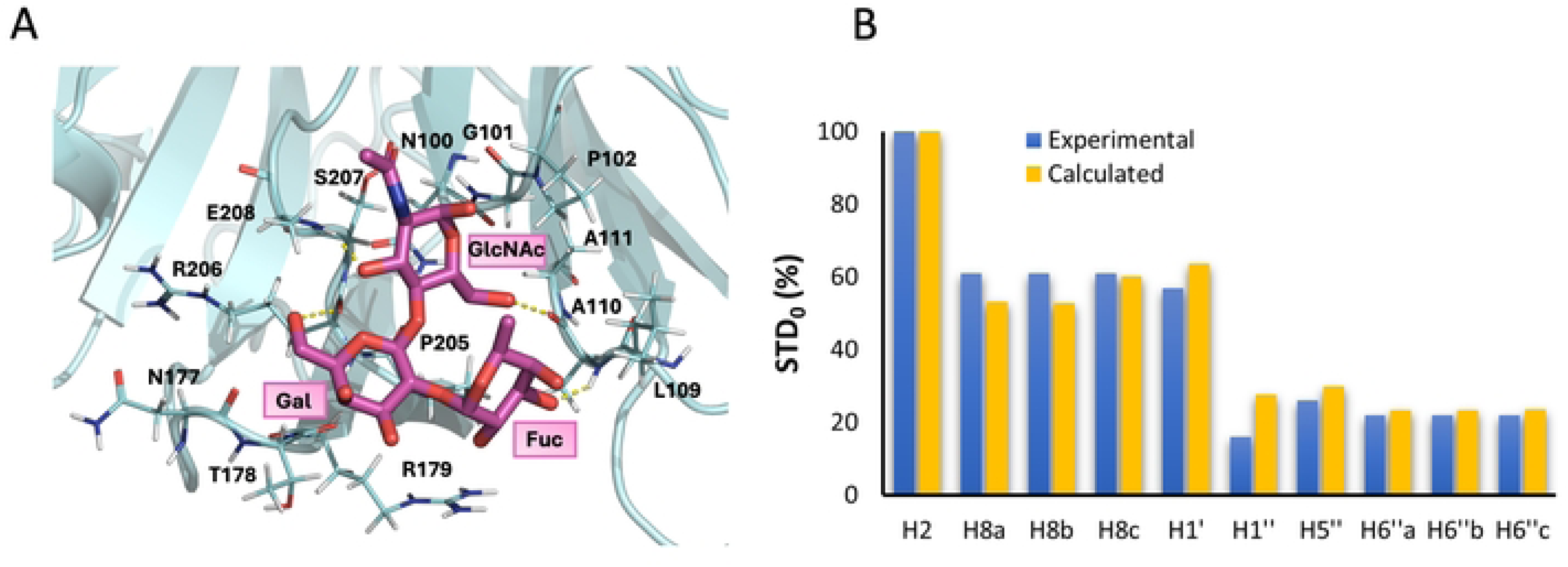
**A** Proposed NMR-validated 3D model for the binding of bovine RVC VP8* and H2 antigen. In magenta is shown H2 antigen with α configuration. This frame corresponds to frame 321 (the lowest NOE *R*-factor) of the MD simulation (protein in cyan-colored cartoon, ligand in magenta sticks) of the complex. Hydrogen bonds are shown in yellow dashed lines. **B** Comparison between the calculated (yellow bars) and experimental (blue bars) relative STD_0_ factors (binding epitope mapping) of the non-exchangeable protons of the H2 antigen bound to the bovine RVC VP8*. A NOE R-factor of 0.11 was obtained using a cutoff of 12 Å.

On the other hand, when comparing the proposed 3D model with the x-ray structure of the complex between human RVC VP8* and the antigen A trisaccharide (PDB: 5ZHO), a reorientation of the antigen H2 trisaccharide can be observed. This is due to loop βK-βL in human RVC being shorter in the bovine protein (where the loop is called βK-β2 instead), resulting in a modification of the trisaccharide binding site (S6 Fig) according to the sequence and structure analysis mentioned above.

## Discussion

Interaction with cell-surface glycans, particularly HBGAs, has been shown to play a crucial role in the pathogenesis, host tropism, and interspecies transmission of RVA [10,28,35,38]. Multiple genotype-dependent interactions with glycans have been described, such as P[4], P[6], and P[8] binding to H1 antigen [39], P[9], P[14], and P[25] to A antigen [40,41], and P[11] to LNnT and LNT [42]. However, little is known about the relevance of host glycan profiles in infections caused by non-A rotaviruses. Among non-A RV species, RVB and RVC are the most commonly found infecting humans and animals [22,43].

A recent study revealed that human RVC VP8* protein specifically binds to the A antigen HBGA, although interactions with glycans of animal genotypes within this species remain unexplored [26]. Thus, this study aimed to determine the HBGA-binding specificity of bovine P[3] RVC VP8* and to understand its structural basis and the molecular mechanisms behind this interaction. Our results demonstrate for the first time an interaction between bovine RVC VP8* and the H2 antigen (Fuc-α1,2-Gal-β1,4-GlcNAc). This interaction appears to occur through its disaccharide precursor (LacNAc), as determined by STD-NMR, revealing that it involves primarily N-acetylglucosamine, which has previously been identified as a key binding saccharide in other studies [15,44].

The biological relevance of this interaction lies in the fact that bovine milk is a rich source of type II oligosaccharides, including LacNAc [45]. This suggests that the bovine P[3] RVC genotype may have specifically adapted to its host glycans. Similarly, VP8* proteins from bovine and human strains of P[11] RVA are known to bind specifically to LacNAc and LNnT (Galβ1,4-GlcNAcβ1,3-Galβ1,4-Glc) [42,46]. Interestingly, human P[11] RVA VP4 originates from a genetic reassortment with a bovine strain, thereby crossing the species barrier [46]. Additionally, studies have shown that saliva and immature intestines of neonates and young children are rich in poly-LacNAc, explaining the epidemiology of RVA genotype P[11][47].

Regarding RVC, a Brazilian study demonstrated its zoonotic potential based on the phylogenetic sequence of the VP6 gene [48]. Genetic reassortments between human-porcine and bovine-porcine strains, involving the NSP4, NSP5, VP3, and VP6 genes, have also been reported [22,49]. While the potential for interspecies transmission, such as from bovine to human, remains unclear, the ability of bovine RVC to bind LacNAc, found in neonates and young children, raises the possibility of zoonotic transmission.

Despite the general structural similarity of bovine RVC VP8* to galectins, RVA VP8*, and to the human RVC VP8* alone and bound to A trisaccharide, the bovine strain has evolved distinct structural features, including a wider cleft between the β-sheets, additional loops, and β-strands. The wider cleft in bovine VP8* results mainly from three variations, the deletion of three residues in the βK-β2 loop, the presence of a Leu at the end of β1 beginning of the β1-βB loop, and the conformational change of βJ-βK to include a long flexible loop, leading to significant changes in the binding site of the A trisaccharide. These alterations affect glycan-binding mechanisms in bovine P[3] RVC VP8*.

Sequence alignment of VP8* proteins from different RVC genotypes confirmed the notable variations, such as the three-amino acid deletion in the βK-β2 loop and the substitution of Trp with Leu at the end of β1 beginning of β1-βB loop. These changes were observed exclusively in bovine P[3] and porcine P[5] genotypes. These structural and sequence differences may explain the shift in recognition from the A antigen to LacNAc and H2 glycans in bovine P[3] and possibly porcine P[5]. This may also clarify why a recent study found no binding to saliva samples by P[3] and P[5], as these genotypes appear to lack affinity for type I antigens [26].

Similar to P[11] RVA, the specific binding between bovine RVC VP8* and the H2 antigen strongly suggests that H2 may function as a cellular adhesion factor for bovine RVC. Molecular docking studies revealed that the H2-binding site is located in the cleft between the β-sheets, interacting with the βK-β2 loop via residues Arg206 and Glu208. Notably, E208 is one of the least conserved residues among human and animal RVC VP8* sequences. Furthermore, shortening of the βK-β2 loop appeared critical for H2 binding, leading to a reorientation of the ligand to accommodate the binding pocket. This H2 recognition mechanism appears to be specific to RVC, as the interaction between bovine and human P[11] RVA VP8* proteins with LNnT (the core H2 antigen) occurs at the βK-βH cleft. Although the cleft width is similar to that in RVC, the βC-βD loop is shorter, promoting a shift in the binding site (S7 Fig).

Our study reveals that, like RVA, RVC shows a diverse range of interactions with HBGAs, demonstrating evolutionary adaptation to host genetic factors and glycans. This work also raises epidemiological questions about P[3] RVC, such as its prevalence in bovine populations and potential cross-species infections involving sheep or other animals. Their high affinity for LacNAc and H2 antigens suggests that the gastrointestinal epithelium of neonates and young children could serve as a target for infection. Further studies are needed to investigate the global prevalence of P[3] RVC, as well as the HBGA-binding profiles of other genotypes, such as porcine P[5]. Understanding these binding mechanisms will provide valuable insights into zoonotic transmission and species barriers. Finally, the role of VP8* and its ligands in mediating RVC cell entry and replication remains to be further explored.

## Materials and methods

### Expression and purification of RVC VP8* protein

The cDNA encoding VP8* galectin domain (amino acids 64-228) from the VP4 protein of Toyama bovine strain RVC belonging to P[3] genotype (GenBank accession number AB738415) was obtained as synthetic gene cloned into the pMX plasmid provided by GeneART technologies (ThermoFisher) with codon optimization to improve its expression yield in *E.coli*. The sequence included a *Bam*HI cleavage site at the 5’ end and a *Sma*I cleavage site at the 3’ end for their cloning into the pGEX-2T expression vector (GE Healthcare) to express N-terminal GST tag. The recombinant protein GST::VP8* protein was expressed in *E.coli* BL21 (DE3) (Novagen) induced with 0.1 mM isopropyl-β-D-1-thiogalactopyranoside (IPTG) at 18°C at 250 rpm for 18 h. The fusion VP8* protein was purified by affinity chromatography using GSTrap column coupled to an ÄKTA prime FPLC system (GE Heathcare) according to manufacturer protocol. The protein was then concentrated and buffer exchanged to PBS using 10kDa Amicon® tubes (Millipore) and quantified using a fluorescence-based protein quantification detection kit (Quant-iT Fluorometer, Qubit Protein Assay Kit, Q33212, Invitrogen) and concentrated to 5 mg/mL in PBS and stored at -80°C for further use. The PBS composition for STD-NMR experiments was as follows: 10 mM Na_2_HPO_4_, 1.8 mM KH_2_PO_4_, 2.7 mM KCl, and 95 mM NaCl pH 7.4, instead of the standard 137 mM. The sequence is included as a FASTA file in supplementary data (S2 Table).

### Glycan microarray screening

Glycan microarray analysis involved utilizing a synthetic glycan library prepared chemoenzymatically to represent key glycan structures from various origins, such as mammals, invertebrates, and plants [29] (S1 Fig). The fusion GST-VP8* P[3] bovine RVC protein was applied to individual glycan microarray at 250 µg/ml concentration in binding buffer (25 mM Tris, 150 mM NaCl pH 7.5, 2 mM CaCl_2_/MgCl_2_, 0.01% Tween 20) at room temperature for 5 h. The bound GST-VP8* protein was detected using a mouse anti-GST tag monoclonal primary antibody (1:2,000) (Thermo Fisher Scientific, MA4-004) and Alexa Fluor^TM^ 647 rabbit anti-mouse IgG (H+L) (Thermo Fisher Scientific, A21239) secondary antibody (1:1,000). A solution of GST (250 µg/ml) was included as a negative control. The slide was washed with binding buffer and water. Relative fluorescence units (RFU) were measured with the Agilent G265BA microarray scanner system and ProScanArray^®^ Express software from Perkin Elmer was used for image analysis. RFU with local background subtraction from four replicate spots were averaged and represented with GraphPad Prism in the form of histograms along with the standard deviation. The glycan microarray library is illustrated in S1 Fig.

### VP8* glycan binding assays

To perform these assays, a glycan panel, comprising biotinylated HBGAs (Le^a^, Le^b^, LacNAc, lacto-N-biose (LNB), H type-1 (H1), H2, A trisaccharide (Atri), B trisaccharide (Btri), Le^x^, Le^y^, sialyl Le^x^ (SLe^x^) and gangliosides (GM3) (GlycoNZ), was utilized (S1 Table). These glycans consist of biotinylated neoglycoconjugates linked to poly (N-2-hydroxyl acrylamide) with a size range of 30 to 50 kDa. F96 black plates with immobilized streptavidin (Nunc), were coated with biotinylated oligosaccharides (2 μg/ml) in deionized water and incubating for 1 h at 37°C as described previously [15]. After plate functionalization, a 0.05% Tween 20 in PBS (PBS-T) wash was conducted, and the plates were blocked with 3% BSA in PBS-T for 1 h at 37°C. Subsequently, the plate was washed three times with PBS-T, and proteins were introduced and incubated at 4°C overnight at a concentration of 10 μg/ml in PBS-T with 0.1% BSA.

The following day, after three washes with PBS-T, a rabbit anti-GST polyclonal antibody (1:2,000) (Sigma-Aldrich) was added, and the plate was incubated for 1 h at 37°C. Subsequently, the plate was washed three times with PBS-T and was incubated for 1 h at 37°C with a horseradish peroxidase (HRP)-conjugated goat anti-rabbit antibody in PBS-T (1:10,000). Following the final three washes, binding was measured using the QuantaBlue reagent kit (ThermoFisher) following the manufacturer’s recommendations. Fluorescence units were determined using a Fluoroskan microplate reader (ThermoFisher). Each binding assay was performed in triplicate. GST was used as negative control and P[14] RVA GST-VP8* [40] was used as positive binding control, the amino acid sequence is included as FASTA file (S2 Table).

### STD-NMR experiments

All the STD NMR experiments were carried out in PBS D_2_O buffer, pH 7.4. The protein concentration was 50 μM and the ligand concentration was 2 mM. STD NMR spectra were acquired on a Bruker Avance III 700.25 MHz at 278 K. The on- and off-resonance spectra were acquired using a train of 50 ms Gaussian selective saturation pulses using a variable saturation time from 0.5 s to 6 s, and a relaxation delay (D1) of 6 s. The residual protein resonances were filtered using a T1ρ-filter of 10 ms. All the spectra were acquired with a spectral width of 9 kHz and 24K data points using 512 scans in saturation times of 0.5 and 0.75 s; 256 scans in 1, 1.5 and 2 s; and 128 scans in 3, 4, 5 and 6 s. The *off-resonance* frequency was set to 40 ppm in all cases, while the *on-resonance* spectra were acquired by selective saturation of protein aliphatic using an irradiation frequency of -2 ppm to avoid direct irradiation to the ligand. To get accurate structural information from the STD NMR data and to minimize any T1 relaxation bias, the STD build-up curves were fitted to the mono-exponential equation STD(t_sat_) = STD_max_*(1-exp(-k_sat_*t_sat_)) (S8 Fig), calculating the initial growth rate STD_0_ factor as STD_max_*k_sat_ and then normalizing all of them to the highest value [50].

### Protein preparation

For the crystallization of the VP8* protein from the bovine RVC P[3] genotype, thrombin (GE Healthcare) was employed to cleave the GST tag. Digestion of one milligram of each GST-VP8* sample was carried out with 10 units of thrombin at 28°C for 31 hours. Subsequently, an affinity chromatography step was performed on a GSTrap column (GE Healthcare). In this procedure, cleaved GST or VP8* that had not been sufficiently cleaved and remained fused to GST were retained on the column, while the digested VP8* successfully passed through.

Following the affinity chromatography, gel filtration chromatography was carried out using a Superdex® 200 column coupled to an Äkta avant chromatography system to eliminate aggregated protein and residual thrombin used for cleavage. Finally, protein concentration was achieved using Amicon® Ultra (0.5 ml) 10kDa concentration tubes (Millipore), following the manufacturer’s specifications. Quantification was performed using the Qubit Protein Assay Kit, as described above. The protein was then adjusted to a concentration of 10 mg/ml in a solution containing 50 mM Tris, 150 mM NaCl, pH 8.0, in preparation for the subsequent crystallization process.

### Crystallization and data acquisition

Crystals of VP8* were obtained using the sitting drop vapour diffusion technique. Crystallization was achieved by mixing 0.3 μl of a solution containing 10 mg/ml of protein with 0.3 μl of two different screening solutions (JBScreen Classic HTS I and HTS II, Jena Bioscience). Crystals were grown in a 1.5 M Sodium Citrate pH 6.5 mother solution. For diffraction, crystals were cryopreserved by passing them briefly through the mother solution containing 12% (v/v) ethylene glycol. Then, crystals diffracted X-rays to 3 Å resolution, and data collection was conducted in the BL13-XALOC of Alba Synchrotron (Cerdanyola del Vallès, Spain). Subsequently, data integration and reduction were performed with XDS [51] and Aimless from the CCP4 suite [52]. Molecular replacement was conducted with Phaser [53] using an Alphafold2 [54] predicted structure as a template. The definitive structural models were obtained by iterative cycles of tracing with Coot [55] and refining with Refmac [56]. Data collection and refinement statistics are included in Table 1. The Ramachandran plot for refined VP8* showed 95.08% residues in favored regions, 4.92 % in allowed regions, and 0% in outliers.

### Molecular docking calculations

Before running docking calculations, an exhaustive conformational study of the apo protein was undertaken. A classical MD simulation of the apo protein was performed, and the conformational states of the protein were subsequently analysed. The crystal structure of bovine RVC VP8* (PDB: 9EN4) was used as starting coordinates. Protein conformations most frequently observed during the MD trajectory were extracted, imported into the Maestro module of the Schrödinger software, and prepared with the Protein Preparation Wizard [57]. The module PROPKA was employed to predict the protonation state of polar sidechains at pH 7.5 [58]. The hydrogen-bonding network was automatically optimized by sampling asparagine, glutamine, and histidine rotamers. The model was then minimized using the OPLS3 force field [59] and a heavy atom convergence threshold of 0.3 Å. Conformers of the H2 antigen were generated in MacroModel [60] using the MC/SD tool, and 100 different conformers were obtained. Clustering of conformers was carried by heavy atom RMSD to eliminate redundant poses, and 10 clusters were obtained. From each cluster, the lowest energy conformer was chosen based on the potential energy-OPLS3e term. Docking of the different conformers of the H2 antigen to B-RVC VP8* was then performed using Glide [61]. A cubic grid was generated centered on the position of the ligand of the x-ray structure with PDB 5ZHO (complex between H-RVC VP8* and type A antigen), with an outer box length of 20 Å and an inner box length of 10 Å. All ligand conformers were subjected to rigid docking (i.e. protein residues are kept fixed in the initial conformation) using the SP algorithm, obtaining 100 poses for each conformer. Docking poses were then clustered by heavy atom RMSD, and the pose closer to the centroid of each cluster was selected. Finally, selected poses ware assessed against experimental STD NMR data using RedMat and the models with the lowest R-NOE factors were chosen.

### Molecular dynamics calculations

#### Input preparation and equilibration

The initial coordinates of the B-RVC VP8*-H2 antigen complex and of the apo-B-RVC VP8* were built from the coordinates of the potential candidates obtained from docking simulations. The MD simulation setup and equilibration were performed with the BioExcel Building Blocks (*BioBB*) library [62]. The ligands were parametrized and minimized using the *acpype* and *babel* modules, respectively, of BioBB (*biobb_chemistry.acpype* and *biobb_chemistry.babel*). The minimization of the ligands was performed with the steepest descent method and the GAFF force field. The topologies of the complexes were generated using the *biobb_amber.leap* module, and the ff14SB [63] and GAFF [64] force fields were used to parametrize [63] protein [64] and ligand, respectively. Subsequently, the *biobb_amber.sander* module was employed to minimize, firstly, the protein protons using positional restraints of 50 kcal/mol·Å^2^ on the protein heavy atoms and, secondly, the whole protein structure using positional restraints of 500 kcal/mol·Å^2^ on the ligand to avoid potential changes in ligand orientation due to protein repulsion. Then, each protein-ligand complex was immersed in a TIP3P [65] truncated octahedron water box with a distance from the protein to the box edge of 9.0 Å and Periodic Boundary Conditions, followed by the addition of a 150 mM concentration of NaCl. Each solvated system was minimized using the steepest descent protocol and applying positional restraints of 15 kcal/mol·Å^2^ to the ligand, followed by heating up to 300 K over 2500 steps applying the Langevin thermostat [66] with a collision frequency of 1 ps^−1^ and positional restraints on the ligand of 10 kcal/mol·Å^2^ (for this, the *biobb_amber.sander* module was used). Next, each system was subjected to NVT followed by NPT equilibration of 100 ps each. A non-bonded interactions cutoff of 10.0 Å, the SHAKE algorithm for constraining the length of bonds involving hydrogen atoms, the Langevin thermostat with a collision frequency of 5 ps^−1^, and smooth positional restraints on the ligand (5 and 2.5 kcal/mol·Å^2^ for NVT and NPT, respectively) were employed. During the NPT equilibration, a pressure of 1 bar was kept constant using isotropic position scaling with a pressure relaxation time of 2 ps.

#### MD simulation

A 500 ns of MD production run was carried out for each complex on a AMD-Ryzen 4xGPU 3070 Computing Cluster using the *pmemd.cuda* module of AMBER 20 [67]. The production dynamics was performed at a constant temperature of 300 K, by applying the Langevin thermostat [66] with a collision frequency of 1 ps^−1^, and a constant pressure of 1 bar (using isotropic position scaling with a pressure relaxation time of 1 ps). A non-bonded interactions cutoff of 9.0 Å, periodic boundary conditions (PBC) [68], and the Particle Mesh Ewald method [69] (PME) to account for the long-range electrostatic effect were employed. The SHAKE algorithm [70,71] was also employed, thus allowing 2 fs between time steps. Trajectory coordinates were saved every nanosecond.

The conformational space sampled by the apo-protein during the trajectory was clustered using the DBSCAN algorithm implemented in AMBER, obtaining 10 clusters representative of the conformations adopted by the protein during the simulation.

The analysis of the MD trajectories was performed using the *cpptraj* module (version *4.25.6*) of AMBER 20 [67]. The evolution of protein and ligand RMSD over the simulation time was calculated against the first frame of the trajectory. To monitor ligand orientation and dynamics within the protein binding site, MD trajectories were aligned based on the protein backbone atoms within 5 Å of the ligand (in the first frame) and, subsequently, the ligand backbone RMSD was calculated in-place (no superposition).

### RedMat calculations

RedMat calculations were performed, using a rotational correlation time of the protein of 11.4 ns, as calculated with the HYDRONMR package [72] through the NMRbox shared resource for NMR software [73], we selected irradiated atoms in methyl protons and a dissociation constant of 500 μM. The ligand and protein concentrations were 2000 μM and 50 μM, respectively, according to the experimental conditions. The protons that are averaged in the experimental binding epitope were not considered in the calculation as they do not represent the real individual values.

### Statistical analysis

To analyze statistical differences in the ELISA-like glycan binding experiments student’s t-test was used considering statistically significant p-value< 0.05. Statistical analyses were performed with GraphPad Prism version 5.0 for Windows (GraphPad Software).

### Alignment analysis

Reference sequences from different RVC genotypes were obtained from GenBank. Sequences were aligned with Clustal omega [74] and the amino acids conservation was analyzed using PRALINE multiple sequence alignment [75].

## Data availability

The X-ray crystallographic coordinates reported for the structure of VP8* from the bovine RVC P[3] genotype has been deposited at the Protein Data Bank with the accession code 9EN4.

## Acknowledgments

This work was supported by a grant from the Spanish Ministry of Science and Innovation, Carlos III Health Institute (grant PI20/00801), and by a research grant to N.N.-L. from the Conselleria d’Educació, Cultura i Esports, Generalitat Valenciana (grant ACIF/2020/076). N-C. Reichardt and S. Serna acknowledge funding by grant PID2023-153181OB-I00 from AEI/ 10.13039/501100011033 and Gobierno Vasco BIOCART. This research was supported by grants PID2019-110630GB-I00 and PID2022-141621NB-I00 to P.C. funded by MCIN/AEI/10.13039/501100011033 and by “ERDF A way of making Europe”. We would like to thank the IBV-CSIC Crystallogenesis Facility for protein crystallization screenings. The X-ray diffraction data reported in this work were collected in experiments performed at BL13-XALOC at ALBA Synchrotron (Cerdanyola del Vallès, Spain). We thank the local contacts and staff of beamlines for assisting with data collection.

## Supporting information captions

**S1 Fig. Glycan microarray consisting of 155 different glycan structures; A) Glycan structures included on the microarrays. B) Nature of the glycosidic linkages of the N-glycan structures on the microarrays.**

(PPTX)

**S2 Fig. Structural comparison of bovine RVC VP8*, human RVA VP8* and Galectin-3**. A) Representation of β-sheet2 found in bovine RVC VP8* formed by six strands (βC, βD, βG, βH and two small strands β1 and β2) compared to the S-face of Galectin-3 bound to a carbohydrate (PDB: 6KXA) and to a similar face found in human RVA VP8* (PDB: 6H9W). In VP8* the S-face like is blocked by a β-hairpin. B) Representation of β-sheet1 found in bovine RVC VP8* formed by six strands (βA, βL, βB, βI, βJ and βK) compared to the F-face of Galectin-3 (PDB: 6KXA) and to a similar surface in human RVA VP8* (PDB: 6H9W).

(PPTX)

**S3 Fig. Differences in the width of the cleft between bovine RVC VP8* and several AlphaFold (AF) models of VP8* genotypes promoted by the presence of Leu or Trp in loop β1-βB.** Structural comparison of the crystal structure of bovine RVC VP8*, which corresponds to Bov P[3] Toyama, with AF models of genotypes: Por P[1] Cowden, Hu P[2] Bristol, Por P[4] Tochigui, Por P[5] RV0143, Bov P[10] Y-1-04, Can P[11] 174 and Por P[12] 07-109-12. The presence of Leu in Bov P[3] and Por P[5] is shown compared to Trp in the rest of the AF models. Also, the width of the cleft, influenced by the presence of Trp or Leu, is indicated by dashed lines, being wider in the latter. Bov: bovine, Can: canine, Hu: human and Por: porcine.

(PPTX)

**S4 Fig. Study of the dynamics and stability of the MD simulation by RMSD and the corresponding validation using RedMat between Fucα1-2Galβ1-4GlcNAcβ and RVC VP8*.** A) Evolution of the root mean squared deviation (RMSD) of the Fucα1-2Galβ1-4GlcNAcβ (all atoms except the protons considered) with respect to the protein binding site (residues within 5 Å from the ligand). B) Evolution of the NOE *R*-factor of ligand over the 500 ns MD simulation.

(PPTX)

**S5 Fig. Study of the dynamics and stability of the MD simulation by RMSD and the corresponding validation using RedMat between Fucα1-2Galβ1-4GlcNAcα and RVC VP8*.** A) Evolution of the root mean squared deviation (RMSD) of the Fucα1-2Galβ1-4GlcNAcα (all atoms except the protons considered) with respect to the protein binding site (residues within 5 Å from the ligand). B) Evolution of the NOE *R*-factor of ligand over the 500 ns MD simulation.

(PPTX)

**S6 Fig. Superposition of the 3D model extracted from the MD simulation (frame 321) of bovine RVC VP8* and H2 antigen (protein in green-colored cartoon, ligand in orange sticks) with the x-ray structure (PDB: 5ZHO) of human RVC VP8* and A trisaccharide antigen (protein in salmon-colored cartoon, ligand in cyan sticks).**

(PPTX)

**S7 Fig. Structural comparison of bovine RVC VP8* with bovine and human P**[11] **RVA VP8*.** Superposition of the crystal structure of bovine RVC VP8*, which corresponds to Bov P[3] Toyama, with human RVA VP8* P[11] in the absence (PDB: 4YGW) and presence of LNnT (PDB: 4YG0) and with bovine RVA VP8* P[11] in the absence (PDB: 4YG3) and presence of LNnT (PDB: 4YG6). The superposed structures are shown separated for clarity. The Leu in loop β1-βB is shown in all structures, as well as the width of the cleft indicated by dashed lines. The presence of LNnT is also shown as sticks.

(PPTX)

**S8 Fig. STD NMR study of the binding of H2 antigen to B-RVC VP8* with protein irradiation in the aliphatic spectral region.** STD NMR build-up curves for Fucα1-2Galβ1-4GlcNAcα and Fucα1-2Galβ1-4GlcNAcβ, in complex with VP8*. Temperature 5°C. Saturation frequency set at -2 ppm.

(PPTX)

**S1 Table. Biotynilated glycans used in this study. All biotinylated glycan were acquired from GlycoNZ.**

(PPTX)

**S2 Table. Table sequence of bovine RVC VP8* and human P**[14] **RVA in fasta format.**

(PPTX)

**S1 Video. Superposition of the MD trajectories of B-RVC VP8* and H2 antigen in α configuration (protein in cyan-colored cartoon, ligand in magenta sticks) and β configuration (protein in cyan-colored cartoon, ligand in green sticks).**

(MP4)

## Notes

### Competing Interest Statement

The authors have declared no competing interest.

